## Supplementary Figures and Tables for "Iso-orientation bias of layer 2/3 connections: the unifying mechanism of spontaneous, visually and optogenetically driven V1 dynamics"

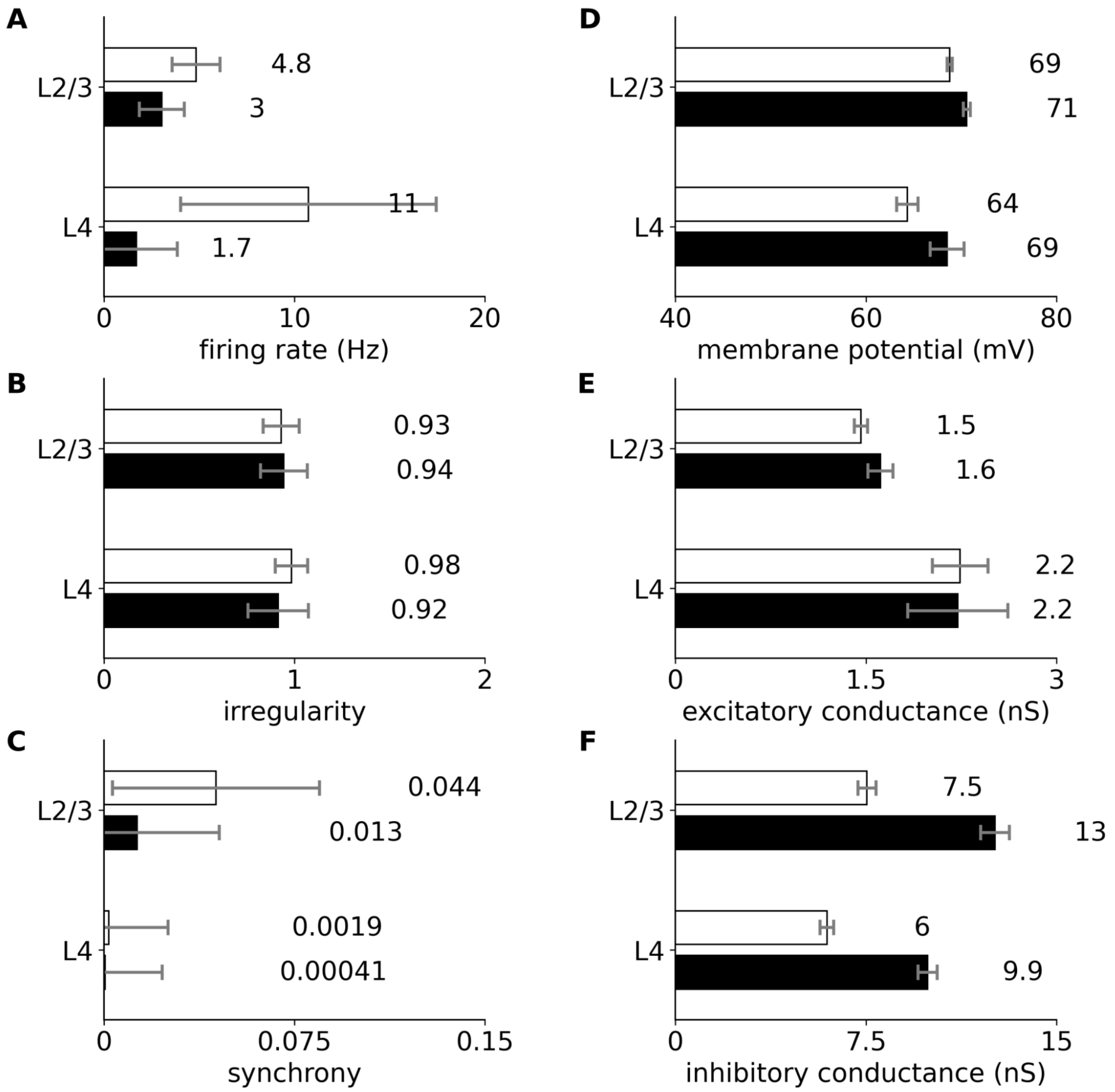


**Figure S1: Statistics of the spontaneous activity of the model.** Six statistics for 40s of spontaneous activity for excitatory (black bars) and inhibitory neurons (white bars) in layer 2/3 and layer 4, with error bars representing the standard error of the mean. The activity of the model in spontaneous state is low and exhibits irregular-asynchronous dynamics, membrane potential resting state of about 70mV and overall realistic subthreshold signals. Refer to Antolík et al. 2024 (Fig. 4A-F) for an extensive description of these analyses and for comparison to the previous iteration of the model (*1*). **(A)** Average firing rate. **(B)** Irregularity of the single unit activity computed as the coefficient of variation of the inter-spike-interval of the neurons. **(C)** Synchrony of the activity computed as the mean correlation between pairs of the spike histograms (bin size of 10ms) of the recorded neurons. **(D)** Mean membrane potential. **(E)** Mean excitatory conductance. **(F)** Mean inhibitory conductance.

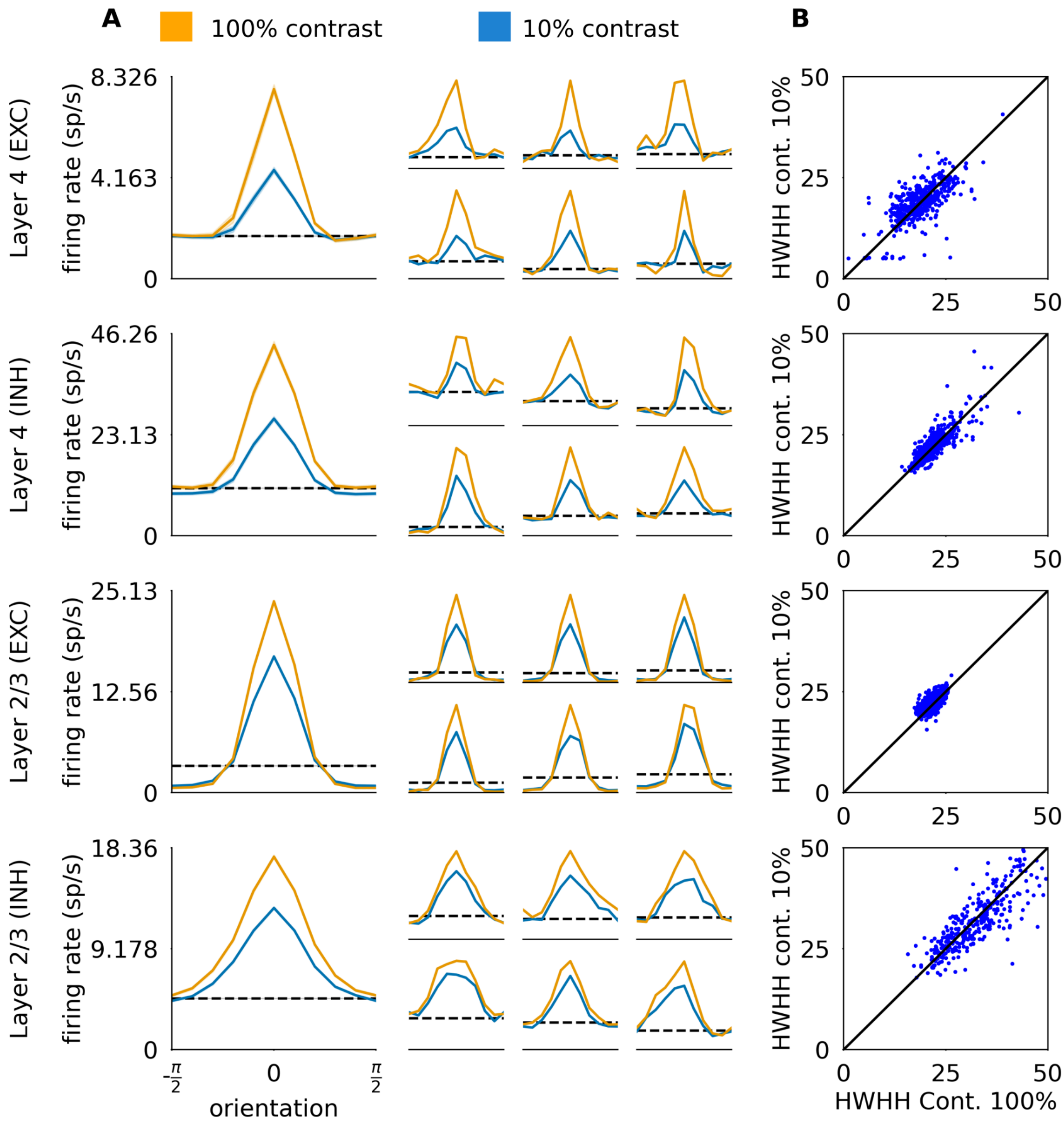


**Figure S2: Orientation tuning properties of the spiking activity of the model.** The orientation tuning curves are computed from the neurons' spiking activity in response to sinusoidal drifting gratings. In all layers, the orientation selectivity of the neurons of the model, measured with the Half-Width at Half-Height (HWHH) of the orientation tuning curves, is contrast-invariant (*2*). Inhibitory neurons remain well tuned with only slightly broader tuning than excitatory ones in line with experimental data (*3*, *4*). Refer to Antolík et al. 2024 (Fig. 6AB) for an extensive description of these analyses and for comparison to the previous iteration of the model (*1*). **(A)** Orientation tuning curves computed based on the mean response of the recorded neurons for 10 orientations of the sinusoidal drifting stimuli for high (yellow line) and low (blue line) contrasts. The baseline levels of activity are marked with dashed lines. Left: Mean orientation tuning curve for the four cortical populations. Right: Single-cell examples. **(B)** Half-width-at-half-weights of the neurons for both low and high contrast stimuli.

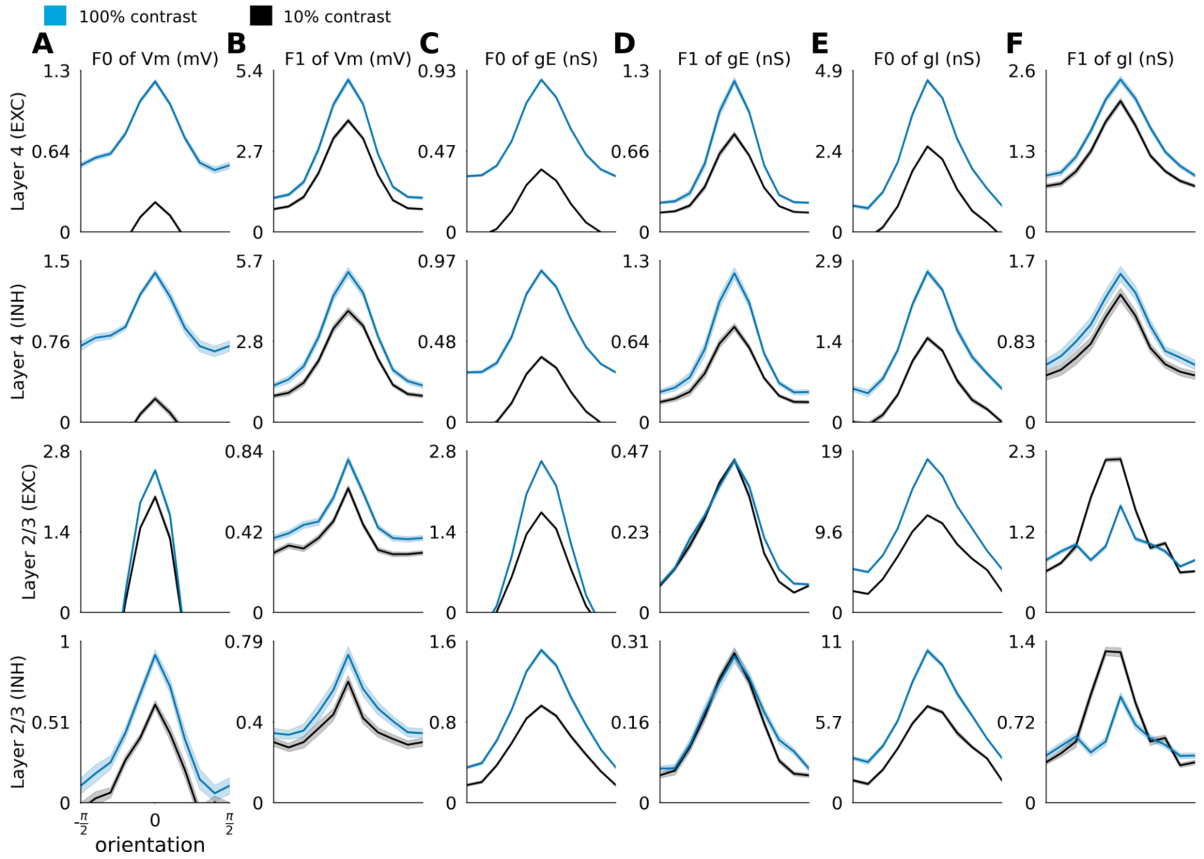


**Figure S3: Orientation tuning properties of the subthreshold signals of the model.** Mean orientation tuning curve for membrane potential and conductances of each of the cortical population for high (blue) and low (black) contrast stimuli. All subthreshold signals exhibit orientation tuning. F0 and F1 refer to the DC and fundamental Fourier components of the neurons' responses to optimally oriented drifting gratings in line with previous intracellular recordings in cat (5). Refer to Antolík et al. 2024 (Fig. 7) for an extensive description of these analyses and for comparison to the previous iteration of the model (*1*). **(A)** F0 component of the membrane potential. **(B)** F1 component of the membrane potential. **(C)** F0 component of the excitatory conductances. **(D)** F1 component of the excitatory conductances. **(E)** F0 component of the inhibitory conductances. **(F)** F1 component of the inhibitory conductances.

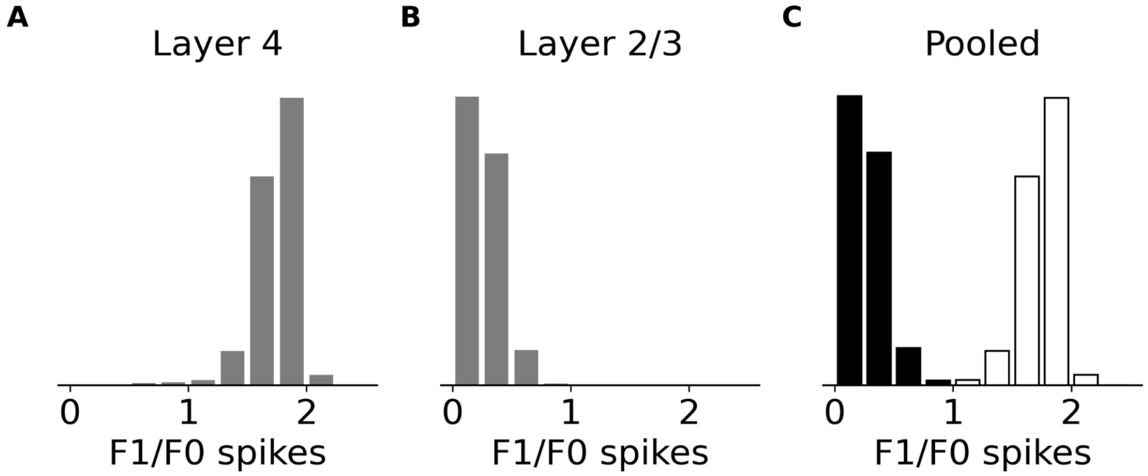


**Figure S4: Modulation ratio of the neurons of the model.** The modulation ratio of the neurons was computed following a drifting sinusoidal gratings protocol. It exhibits a clear bimodal distribution, although with less overlap than observed experimentally in cat and macaque V1 (*6*, *7*). Layer 4 is predominantly populated by simple cells and Layer 2/3 is populated by complex cells. Refer to Antolík et al. 2024 (Fig. 8A) for an extensive description of these analyses and for comparison to the previous iteration of the model (*1*). **(A)** Modulation ratio histogram for excitatory neurons in layer 4. **(B)** Modulation ratio histogram for excitatory neurons in layer 2/3. **(C)** Modulation ratio histogram for excitatory neurons pooled from both cortical layers. Cells identified as complex (F1/F0 < 1) are represented with black bars, cells identified as simple (F1/F0 > 1) are represented with white bars.

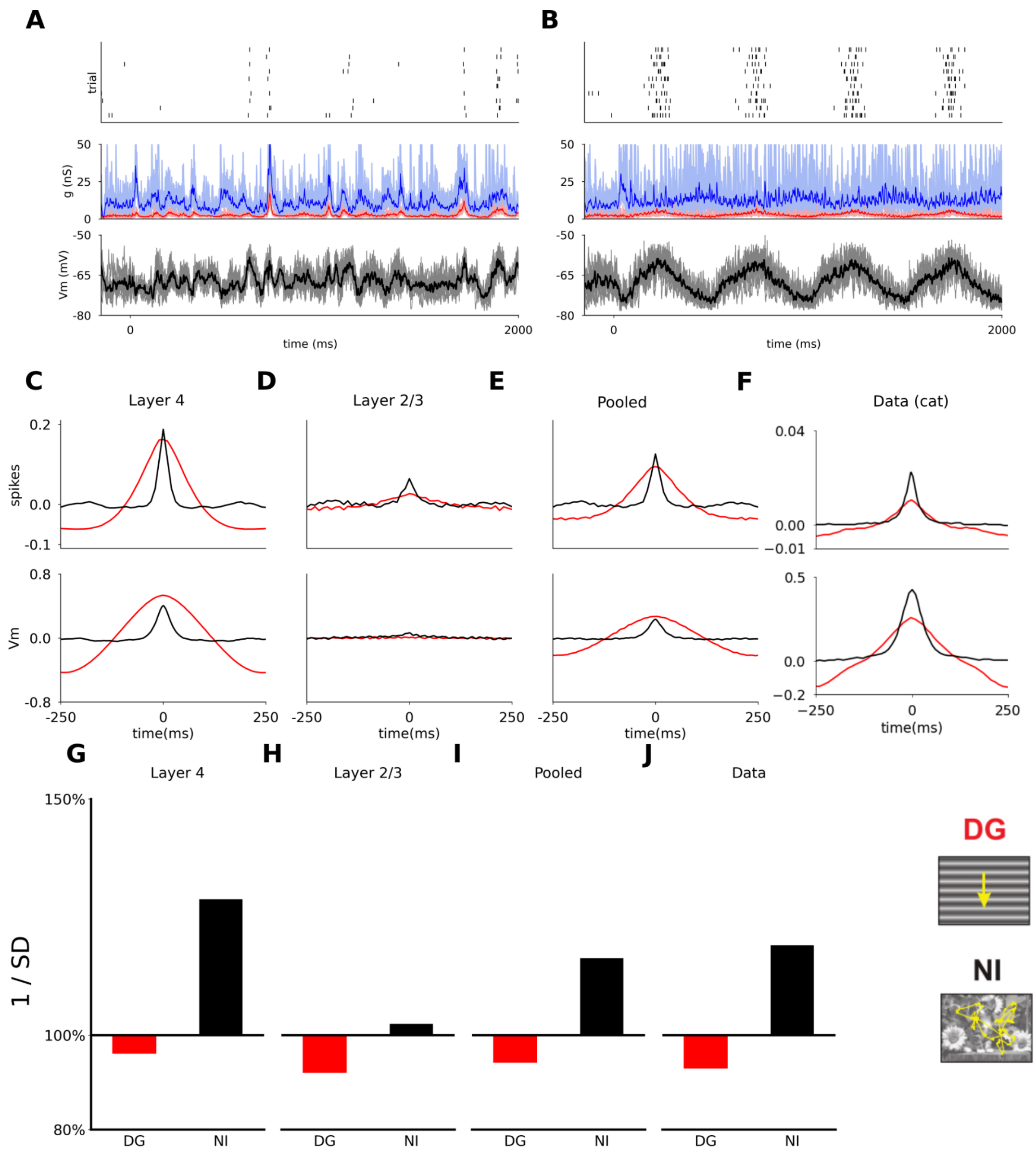


**Figure S5: Trial-to-trial statistic of the response of the model to drifting gratings and natural movie stimuli.** Neurons' responses to a natural movie stimulus with simulated eye movement (NI) exhibit higher reliability and precision than to drifting gratings stimuli (DG). The trial-to-trial cross-correlation response profiles are in good qualitative agreement with the experimental data of Baudot et al. (8), except for layer 4 membrane potential response which exhibit a marginally larger peak for DG than for NI. When pooled across the entire model, the inverse of the standard deviation of the response of neurons relative to spontaneous activity (1/SD) has a profile corresponding to the data of Baudot et al.. Refer to Antolík et al. 2024 (Fig. 10) for an extensive description of these analyses and for comparison to the previous iteration of the model (*1*). **(A)** Response of an example layer 4 excitatory neuron to an NI stimulus. Top panel: raster plot for the ten repetitions. Middle panel: Average excitatory (red) and (blue) inhibitory conductances across trials. Bottom panel: Average membrane potential across trials. **(B)** Same, for a DG at medium contrast (30%). **(C)** Average trial-to-trial cross-correlation for the spike histogram (top) and membrane potential (bottom) for the DG (red) and NI (black) stimuli across all recorded excitatory neurons in layer 4. **(D)** Same, for excitatory neurons in layer 2/3. **(E)** Same, for excitatory neurons pooled across both cortical layers. **(F)** Experimental data in anaesthetised cat. (*8*)**.** **(G)** Inverse of the standard deviation of the response of neurons (1/SD) expressed in percentage relative to spontaneous activity for DG (red) and NI (black) stimuli averaged across all record excitatory neurons in layer 4. **(H)** Same, for excitatory neurons in layer 2/3. **(I)** Same, for excitatory neurons pooled across both cortical layers. **(J)** Experimental data in anaesthetised cat (*8*)**.**

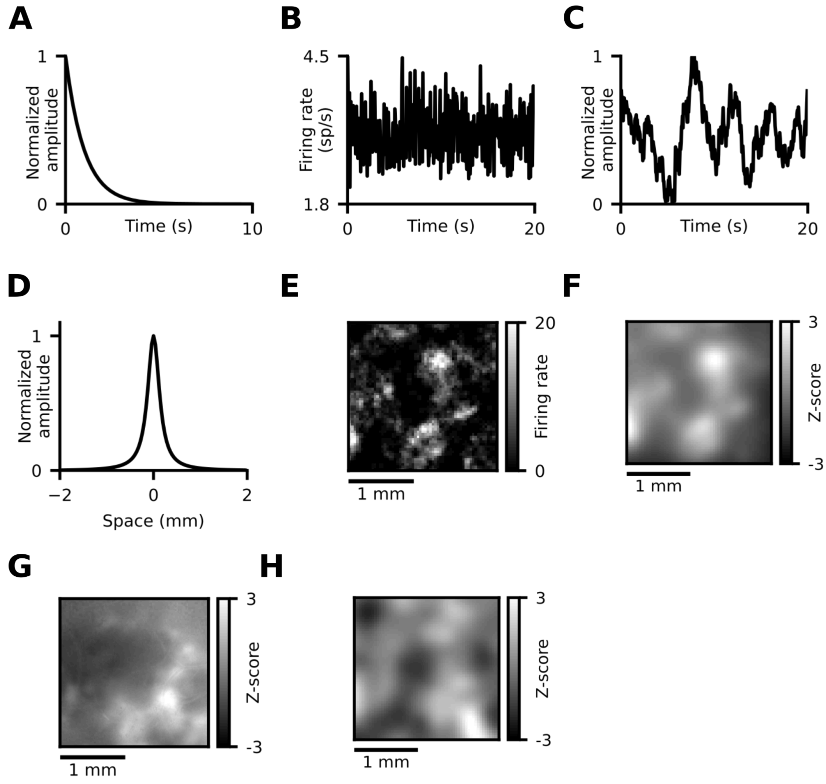


**Figure S6: Constructing the simulated calcium imaging signal. (A)** Experimentally measured (*9*) temporal kernel (exponential decay, $\tau$=1s) of the calcium imaging construct. The rise time constant is ignored as it is much smaller (10ms) than the temporal resolution of our signal (50ms). **(B)** Population mean spiking activity of Layer 2/3 during spontaneous activity. **(C)** Timecourse of the temporally filtered spiking activity, resulting from the convolution with the exponential decay kernel from (A). **(D)** Spatial smoothing kernel derived from a model of light spread within the cortex (*10*), accounting for the diffusion of the spiking signal in the calcium imaging readout **(E)** Example frame of time-binned spiking Layer 2/3 model activity. **(F)** Spiking activity frame from (E), convolved with the light spread kernel from (D). **(G)** Reference frame of ferret cortex activity recorded using calcium imaging (*11*), adapted from Smith et al. (*11*), *Nature Neuroscience*, © 2018 Springer Nature, with permission obtained via RightsLink. **(H)** Example frame of the fully filtered model cortex activity (spatial + temporal), producing a signal visually similar to the experimental reference in (G).

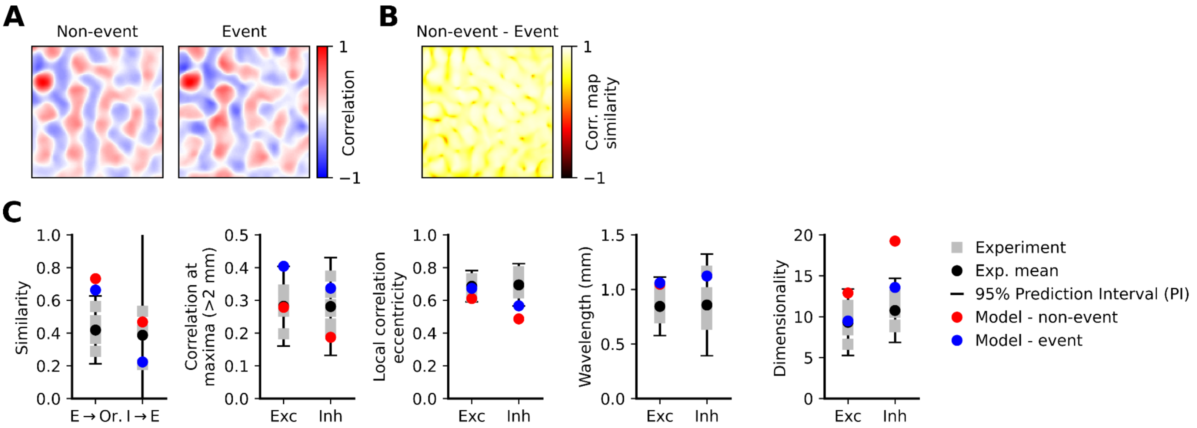


**Figure S7: Comparison of results calculated from entire activity time course vs. only large-scale event maxima. (A)** Spatial correlation maps are similar whether screening for large-scale events is applied or not, on a 120s recording of spontaneous activity data. Screening is done as per Smith et al. (*11*), but with a pixel activation threshold of 2 s.d. and an active frame threshold of 0.8. The decreased pixel activation threshold (from 4-5 s.d. in Smith et al. (*11*)) is due to the lower levels of noise in our model compared to experiment. **(B)** Correlation similarity (see Methods) between non-event- and event-based correlation maps is consistently very high (mean=0.78 ± 0.001), regardless of seed position **(C)** Comparison of metrics measuring the spatial properties of spontaneous activity (see also Figure 3H). Most metrics stay within ferret cortex (*11*, *12*) data ranges, regardless of screening for large-scale events. Omitting screening results in a slight weakening of spatial correlations (still within experimental bounds), and increase in dimensionality (excitatory dim. within, inhibitory dim. outside experimental bounds).

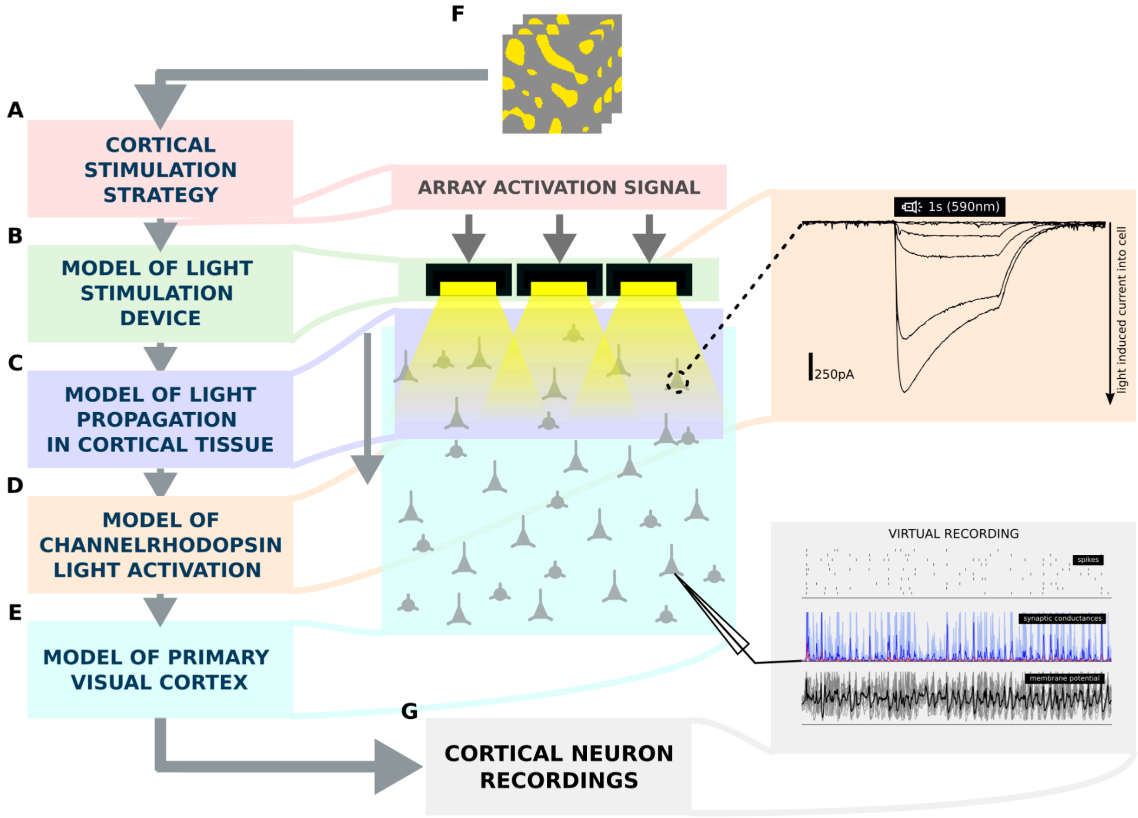


**Figure S8: Detailed schematic of the optogenetic stimulation construct. (A)** the stimulation strategy containing driving signals for a matrix of light emitting elements (MLEE), **(B)** a model of the MLEE **(C)** a model of light propagation through cortical tissue taking into consideration the absorption and diffraction of the light as it travels through the neural substrate. **(D)** a model of channelrhodopsin (ChR) dynamics in transfected cells that transforms a temporal trace of light impinging onto a given cell into a current that is injected into the cell due to the activation of ChR channels, and **(E)** a detailed large-scale spiking conductance based neural model of 5 $mm^2$ of primary visual cortex. Using these 5 simulation elements, we will be able to simulate the activation of cortical population to a specific set of optogenetic stimulation patterns. Figure adapted from by Antolík et al. (*13*) , used under CC BY 4.0.

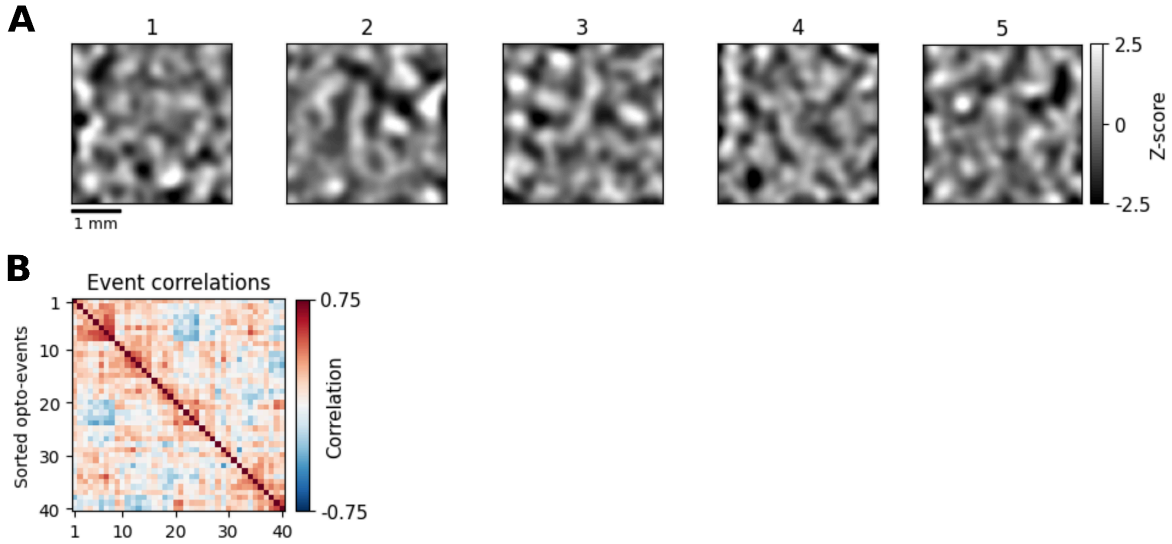


**Figure S9: Model responses to fullfield optogenetic stimulation are variable between trials. (A)** Optogenetically evoked events by fullfield stimulation (defined as the frame of opto-stimulus offset) are distinct **(B)** The sorted spatial correlations between optogenetically evoked events show considerable trial-to-trial variability (inter-stimulus interval = 60s).

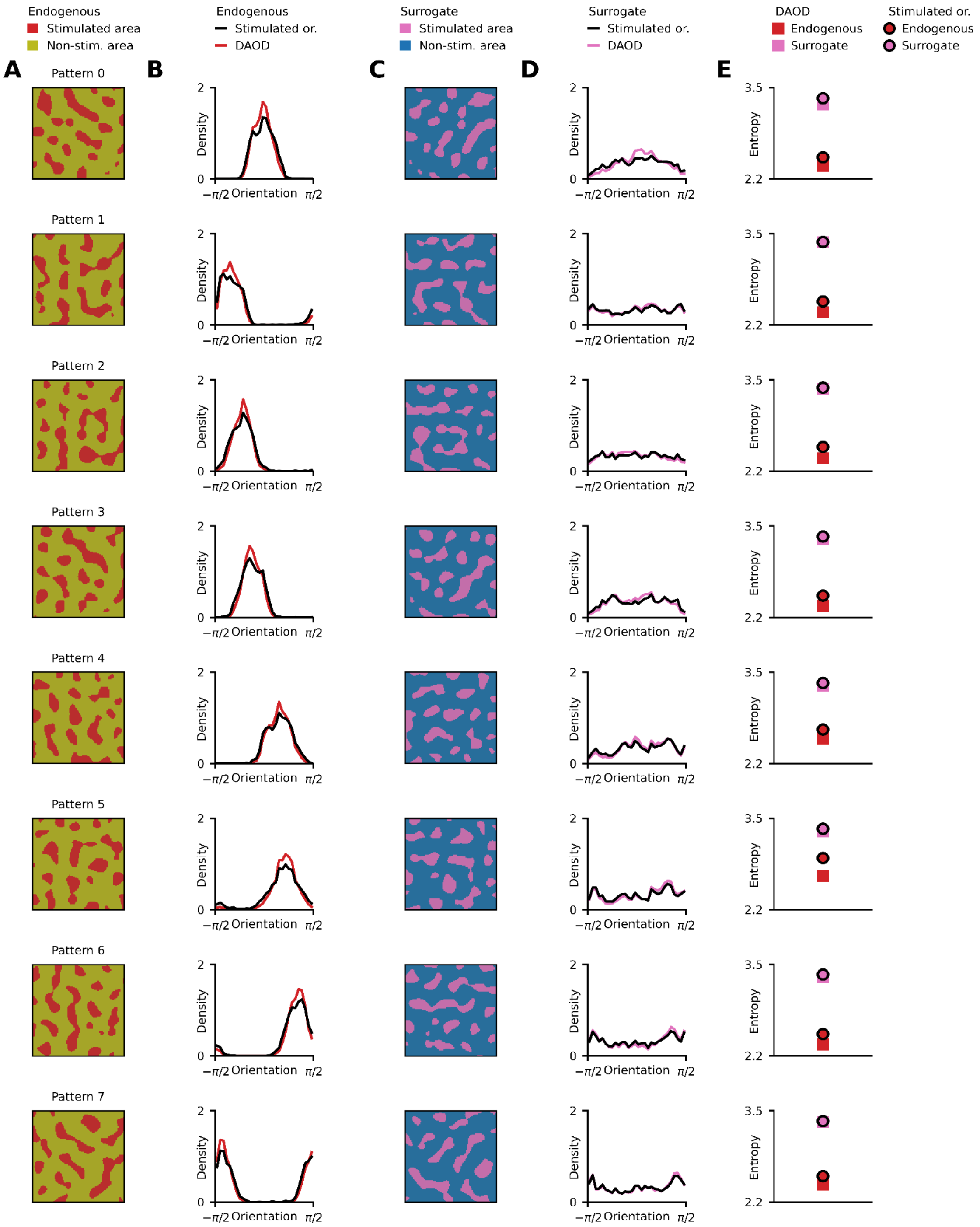


**Figure S10: Distribution of Activity across the Orientation Domain (DAOD) compared to the distribution of stimulated orientations, all stimulation patterns.** **(A)** Individual endogenous stimulation patterns **(B)** For all patterns, the Distribution of Activity across the Orientation Domain (DAOD) under the stimulated area is more sharply oriented than the distribution of orientation preferences (DOP) for the same area, suggesting the selective amplification of the dominant orientation by orientation-biassed connectivity. **(C,D)** Same as (A,B) for surrogate stimulation patterns **(E)** For all patterns, Shannon entropy values of DAODs are lower than corresponding (DOP) entropies, signifying sharper histograms.

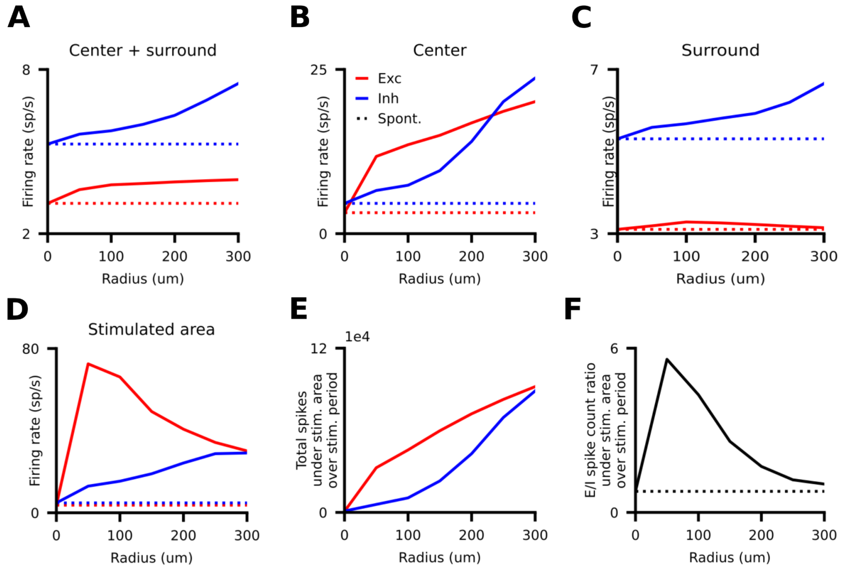


**Figure S11: Firing rates & spike counts during central stimulation. (A)** Mean firing rate of the excitatory and inhibitory subpopulations in the cortex, both continually rising with increasing stimulation radius, inhibitory activity rising faster. **(B)** Mean firing rate in the central area, continually rising as more and more neurons are recruited by stimulation. **(C)** Mean firing rate in the surround, same as Figure 5I **(D)** Mean firing rate under the stimulated area. Although total stimulation input to the cortex stays constant, the stimulation input per stimulated neuron decreases, resulting in a continuous decrease of activity under the stimulated area. **(E)** Although per-neuron excitation decreases with increasing radius, the total number of generated spikes under the stimulated area increases continuously. Thus the local maximum of excitatory activity in the surround cannot be explained by a local maximum in excitatory spikes generated in the centre. **(F)** As a progressively larger area is recruited by central stimulation, the ratio of excitatory/inhibitory spikes generated in the centre is falling continuously towards the spontaneous baseline.

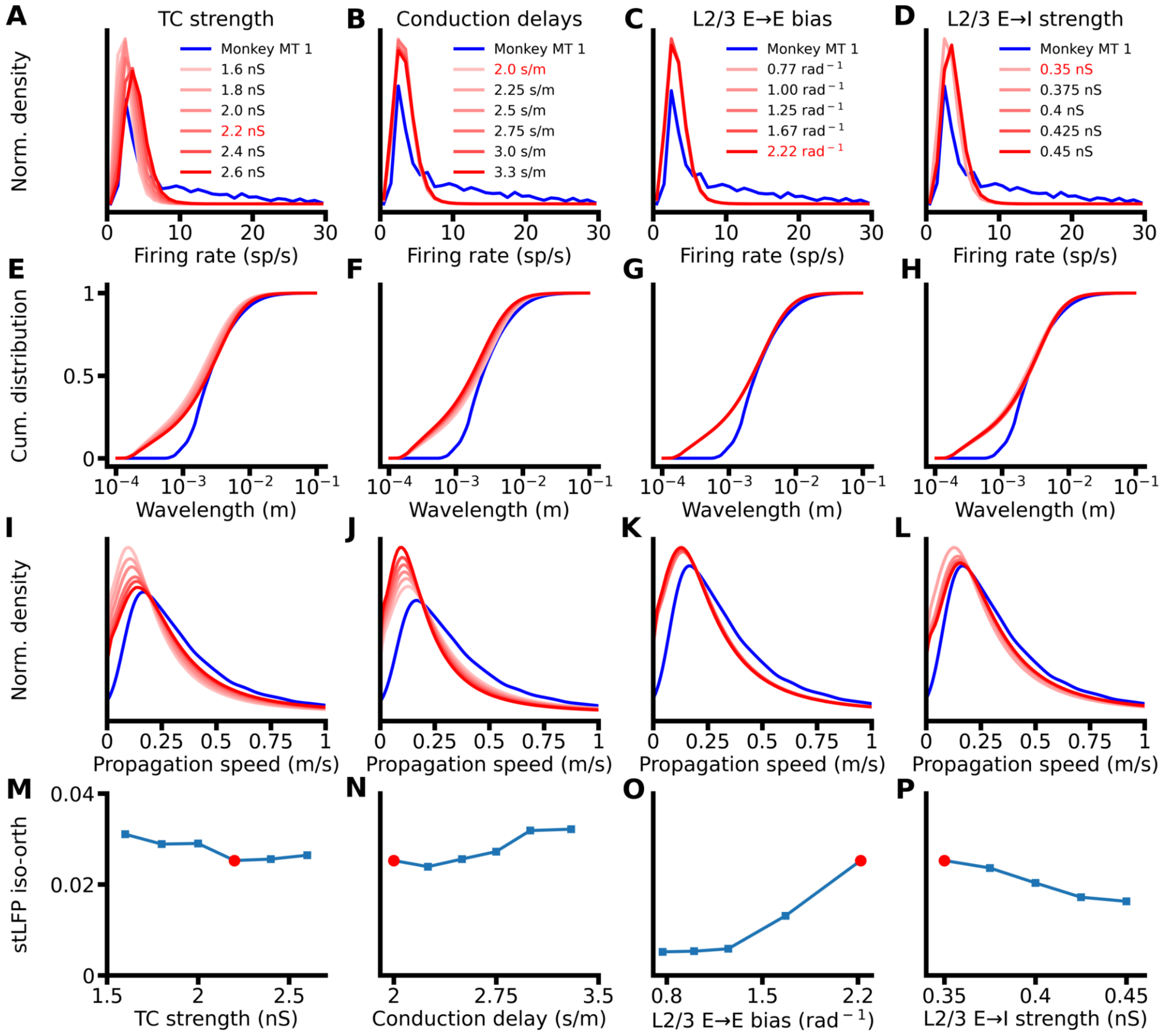


**Figure S12: Impact of the model parameters on STW properties. (A-D)** The distribution of firing rates for six parametrizations varying in TC strength **(A),** six parameterizations varying in conduction delays **(B)**, five parameterizations varying in L2/3 E→E bias **(C)**, and five parameterizations varying in L2/3 E→I strength **(D)** (For detailed choice of parameters see Methods). Red font in legends indicates the parameter values used in the base model. **(E-H)** Same, for the distribution of wavelengths. **(I-L)** Same, for the distribution of propagation speeds. **(M-P)** Same, for the amplitude stLFP residuals difference between recording sites with orientation preference congruent to the reference electrode and sites with orthogonal preference. The red circles indicate the parameter values used in the base model.

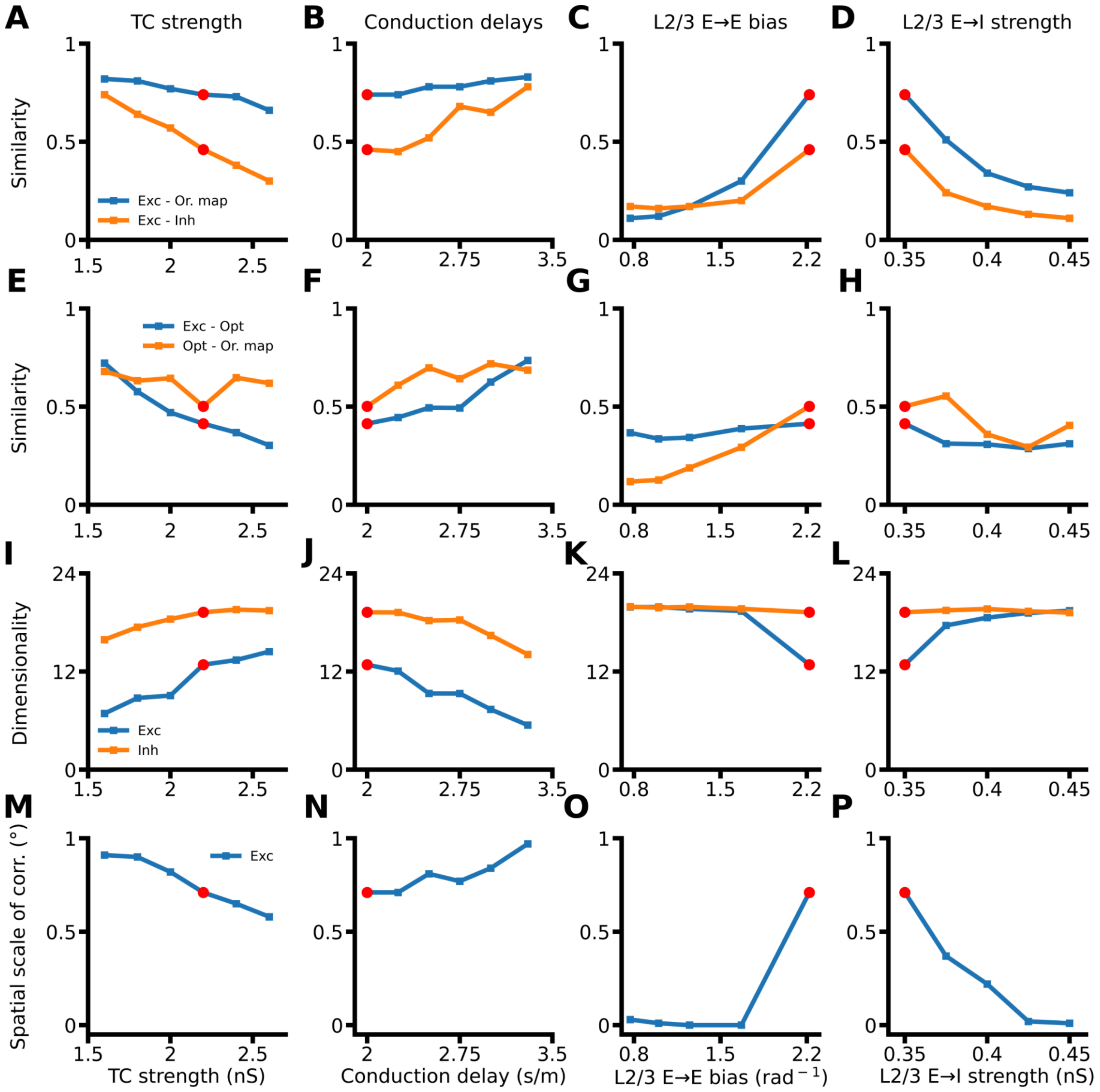


**Figure S13: Effect of the model parameters on the modular activity properties. (A-E)** Similarity between the excitatory CMs and the orientation map (blue), and between the excitatory and inhibitory CMs (orange), for six parameterizations varying in TC strength **(A),** six parameterizations varying in conduction delays **(B)**, five parameterizations varying in L2/3 E→E bias **(C)**, and five parameterizations varying in L2/3 E→I strength **(D)** (see Methods). The red circles indicate the parameter values used in the base model. **(E-H)** Same, for the similarity between the excitatory CMs and the uniform full-field optogenetically evoked excitatory CMs (blue), and between the full-field optogenetically evoked excitatory CMs and the orientation map (orange). **(I-L)** Same, for the dimensionality of the activity of the excitatory (blue) and inhibitory (orange) neurons. **(M-P)** Same, for the spatial scale of correlations.

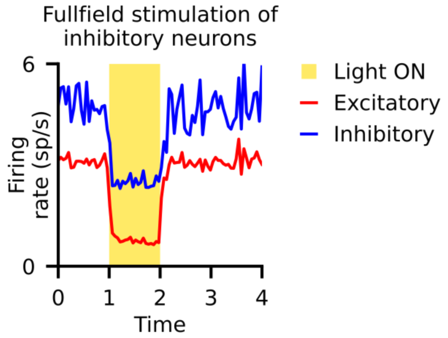


**Figure S14: Full-field stimulation of inhibitory neurons inhibits both inhibitory and excitatory population, in line with dynamical response expected of inhibition-stabilised networks.**

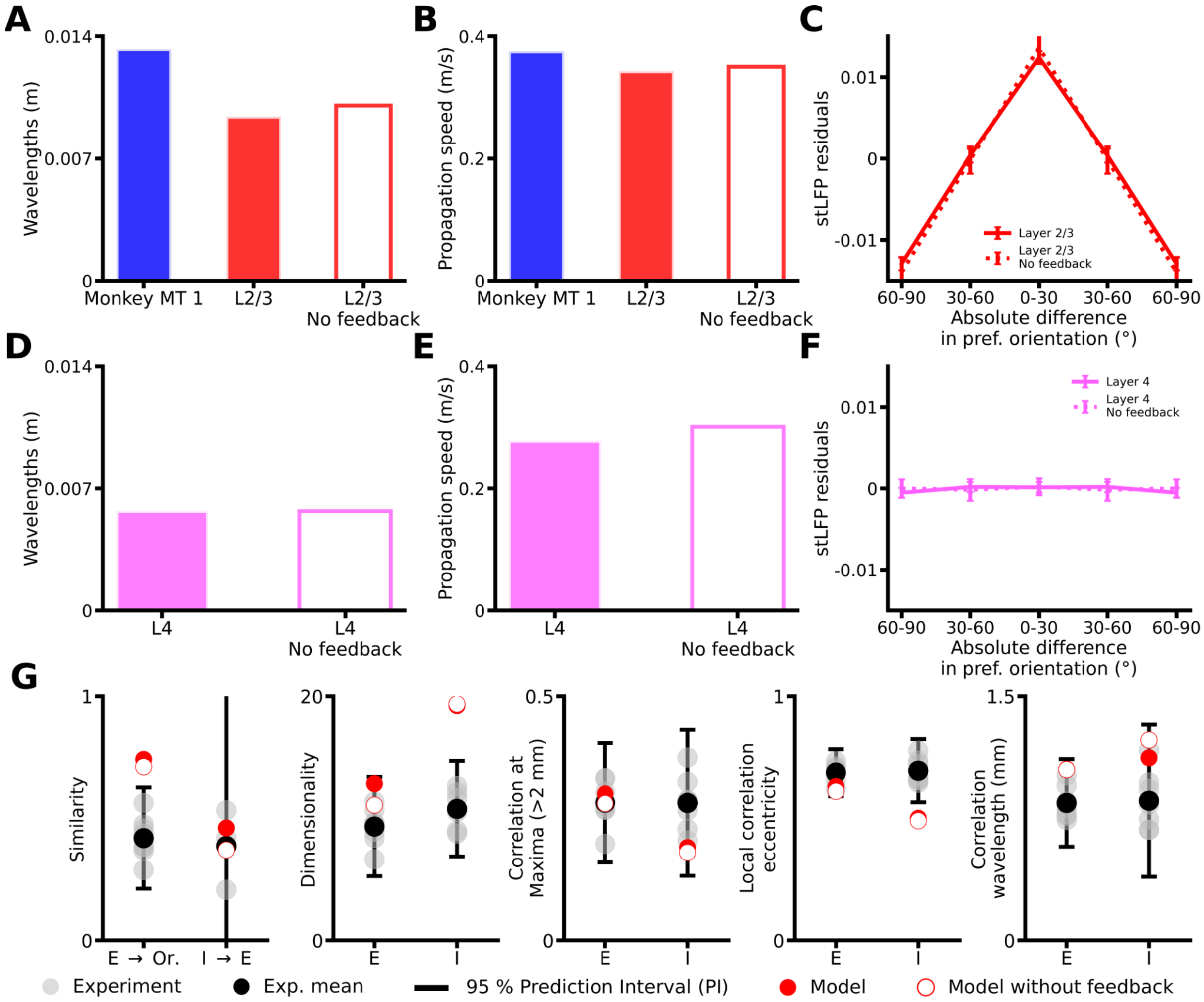


**Figure S15: Effect of the absence of the feedback connectivity on model results. (A)** The average wavelengths in layer 2/3 for the original instance of the model (red, full) and a version where the feedback connectivity is removed (red, hollow), and for the experimental data of Davis et al. for marmoset MT (blue) (*14*), as computed with the Generalized phase analysis. **(B)** Same as (**A)** for the average propagation speeds. **(C)** The average of the spike-triggered LFP residuals in layer 2/3 for the original instance of the model (red, solid) and a version where the feedback connectivity is removed (red, dashed) across reference neurons and recording sites, binned according to their orientation preference differences. Error bars: 95% confidence interval. **(D)** The average wavelengths in layer 4 for the original instance of the model (pink, full) and a version where the feedback connectivity is removed (pink, hollow), as computed with the Generalized phase analysis. **(E)** Same as (**D**) for the average propagation speeds. **(F)** The average of the spike-triggered LFP residuals in layer 4 for the original instance of the model (pink, solid) and a version where the feedback connectivity is removed (pink, dashed) across reference neurons and recording sites, binned according to their orientation preference differences. Error bars: 95% confidence interval. **(G)** Comparison of the layer 2/3 model results with (red, full) and without feedback (red, hollow) for a wide range of metrics also computed in the ferret cortex (gray, with mean represented in black) (*11*, *12*)**.** Experimental data scraped from Smith et al. (*11*) Supplementary Figure 5g, Mulholland et al. (*12*) Figure 4c,e,g,j, 5f.

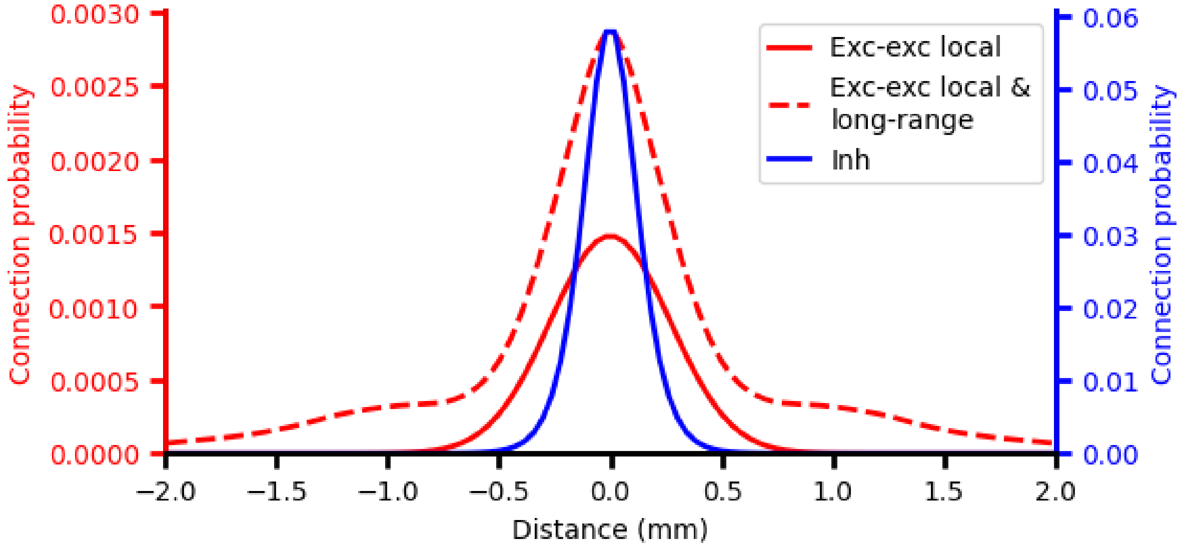


**Figure S16: Spatial extent of excitatory and inhibitory lateral connectivity in model Layer 2/3.** The spatial constants for the excitatory-to-excitatory lateral connectivity in Layer 2/3 are based on the bouton density profiles in Buzás et al. (*17*). All other local connectivity, including the local connectivity of inhibitory neurons, is the result of fitting a hyperbolic distribution to the connection probability profiles in Stepanyats et al. (*18*). The resulting spatial distributions, thus constrained by anatomical data, show a wider local spatial connectivity profile for excitatory neurons than for inhibitory neurons.

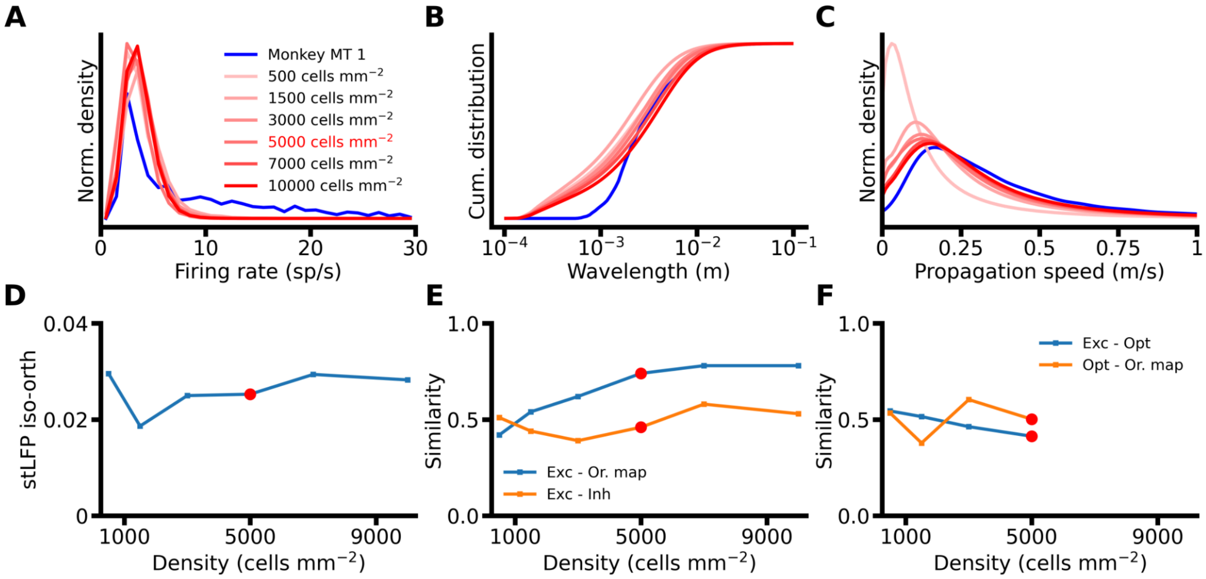


**Figure S17: Effect of cortical density on the model results.** Previous studies have shown that both properties of waves (*14*) and correlation of activity (*15*) are negatively affected by downscaling the neuronal density. The model instance we present here contains 5000 cells per square millimeter which corresponds to a downscaling of roughly one fifth in respect to the real neuronal density in V1 (*16*). This downscaling is necessary for running our computational simulations within a practical amount of time and memory. Several of the metrics we report here are substantially affected by the model neuronal density. When decreasing the density of the model, both distribution of wavelengths and propagation speeds are increased. Moreover, the similarity between spontaneous excitatory correlation maps and the orientation map also decrease as the density of the model is reduced. However, the effect of density considerably saturates once reaching the value of 5000 cells per square millimeter used in this paper. These results therefore indicate that a model needs to have a high enough density to give a quantitatively accurate account of the phenomena that are the focus of this study, but that a downsampling by a factor five or less doesn't significantly affect the results and represents a good compromise between fidelity and computational efficiency. **(A)** The distribution of firing rates for six parametrizations varying in cortical density (see Methods). Red font in legends indicates the parameter values used in the base model. **(B)** Same as **(A)**, for the cumulative distribution of wavelengths. **(C)** Same as **(A)**, for the distribution of propagation speeds. **(D)** stLFP residuals difference between recording sites with orientation preference congruent to the reference electrode and sites with orthogonal preference for six parametrizations varying in cortical density. **(E)** Similarity between the excitatory CMs and the orientation map (blue), and between the excitatory and inhibitory CMs (orange) for six parametrizations varying in cortical density. **(F)** Similarity between the excitatory CMs and the uniform full-field optogenetically evoked excitatory CMs (blue), and between the full-field optogenetically evoked excitatory CMs and the orientation map (orange) for four parametrizations varying in cortical density. We couldn't run these experimental protocols on high density networks due to the large computational cost. The red circles in **(D-F)** indicate the parameter values used in the base model.

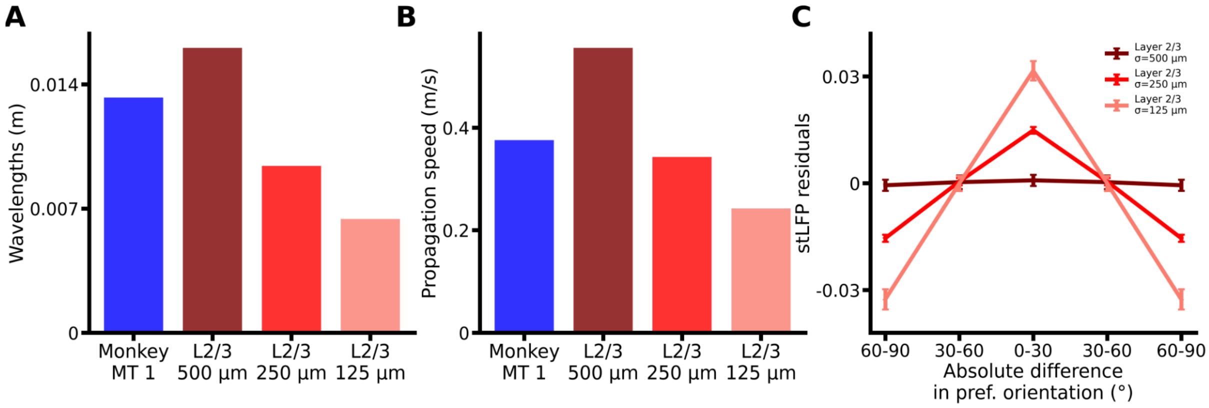


**Figure S18: Dependence of the LFP signal properties on the standard deviation used in the gaussian convolution. (A)** The average wavelengths for model layer 2/3 for standard deviations of 500 μm (brown red), 250 μm (red) and 125 μm (pale red), and for the experimental data of Davis et al. for marmoset MT (blue) (*14*), as computed with the Generalized phase analysis. **(B)** Same as (**A**) for the average propagation speeds. **(C)** The average of the spike-triggered LFP residuals for model layer 2/3 for standard deviations of 500 μm (brown red), 250 μm (red) and 125 μm (pale red) across reference neurons and recording sites, binned according to their orientation preference differences. Error bars: 95% confidence interval.

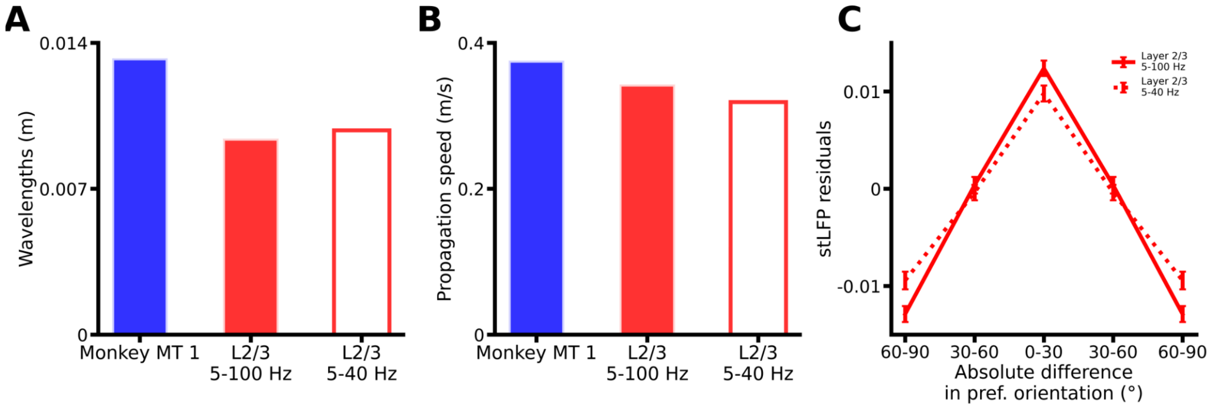


**Figure S19: Dependence of the LFP signal properties on the range of frequencies used in the band-pass filter. (A)** The average wavelengths for model layer 2/3 for band-pass filtering ranges of 5-100 Hz (red, full) and 5-40 Hz (red, hollow), and for the experimental data of Davis et al. for marmoset MT (blue) (*14*), as computed with the Generalized phase analysis. **(B)** Same as (**A**) for the average propagation speeds. **(C)** The average of the spike-triggered LFP residuals for model layer 2/3 for band-pass filtering ranges of 5-100 Hz (red, solid) and 5-40 Hz (red, dashed) across reference neurons and recording sites, binned according to their orientation preference differences. Error bars: 95% confidence interval.

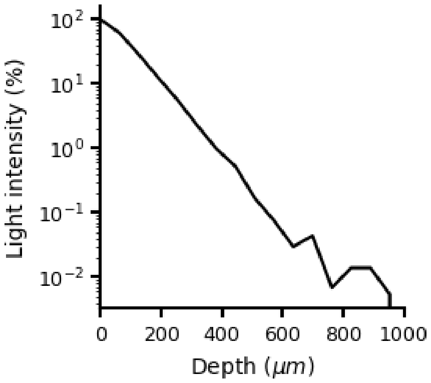


**Figure S20: Exponential light intensity falloff with increasing cortical depth, calculated using the simulations of the 'Human Brain Grey Matter' model implemented in the LightTools software, assuming emitted light at 590 nm.**

| A | Model Summary |
|---|---|
| Populations | Six: V1 Layer 4 (excitatory and inhibitory), V1 Layer 2/3 (excitatory and inhibitory), LGN (ON and OFF) |
| Topology | 2D cartesian space using visual space coordinates (translated for V1 layers to cortical space coordinates through cortical magnification factor) |
| Connectivity | Distant dependent connectivity with functional biases (intra-cortical), Gabor-like (thalamo-cortical) |
| Neuron models | Adaptive exponential integrate-and-fire model (V1), leaky integrated-and-fire model (LGN) |
| Channel models | — |
| Synapse model | Conductance based with exponential decay |
| Plasticity | Short-term plasticity |
| Input | Current fed into the LGN neurons based on the convolution of their center-surround receptive field with the stimulus, and noise. |
| Measurements | Spikes, membrane potential, excitatory and inhibitory conductances |

| B | Populations | | |
|---|---|---|---|
| Name | Elements | Size | Number of neurons |
| LGN ON | Leaky integrate-and-fire | 6°× 6° | 7 200 |
| LGN OFF | Leaky integrate-and-fire | 6°× 6° | 7 200 |
| Layer 4 Exc | Adaptive exponential integrate-and-fire model | 5 mm × 5 mm | 125 000 |
| Layer 4 Inh | Adaptive exponential integrate-and-fire model | 5 mm × 5 mm | 31 250 |
| Layer 2/3 Exc | Adaptive exponential integrate-and-fire model | 5 mm × 5 mm | 125 000 |
| Layer 2/3 Inh | Adaptive exponential integrate-and-fire model | 5 mm × 5 mm | 31 250 |

| C | Connectivity | | | |
|---|---|---|---|---|
| Source | Target | Incomming synapses per post-synaptic neuron | Weight | Description of the connection probabilities between neurons |
| LGN ON | Layer 4 Exc | Uniform within [45,95] interval | 2.2 nS | Gabor distribution |
| LGN OFF | Layer 4 Exc | Uniform within [45,95] interval | 2.2 nS | Gabor distribution |
| LGN ON | Layer 4 Inh | Uniform within [56,84] interval | 2.2 nS | Gabor distribution |
| LGN OFF | Layer 4 Inh | Uniform within [56,84] interval | 2.2 nS | Gabor distribution |
| Layer 4 Exc | Layer 4 Exc | 640 – number of synapses received from the LGN | 0.18 nS | Distance-dependent with 'Push' mechanism |
| Layer 4 Exc | Layer 4 Inh | 384 – number of synpases received from the LGN | 0.35 nS | Distance-dependent with 'Push' mechanism |
| Layer 4 Inh | Layer 4 Exc | 200 | 2 nS | Distance-dependent with 'Pull' mechanism |
| Layer 4 Inh | Layer 4 Inh | 120 | 2 nS | Distance-dependent with 'Pull' mechanism |
| Layer 4 Exc | Layer 2/3 Exc | 404 | 1 nS | Distance-dependent with orientation bias |
| Layer 4 Exc | Layer 2/3 Inh | 242 | 1 nS | Distance-dependent with orientation bias |
| Layer 2/3 Exc | Layer 2/3 Exc | 1435 | 0.18 nS | Local distance-dependent component<br>Long-range distance dependent component with orientation bias |
| Layer 2/3 Exc | Layer 2/3 Inh | 861 | 0.35 nS | Local distance-dependent component<br>Long-range distance dependent component with orientation bias |
| Layer 2/3 Inh | Layer 2/3 Exc | 460 | 2 nS | Distance-dependent |
| Layer 2/3 Inh | Layer 2/3 Inh | 276 | 2 nS | Distance-dependent |
| Layer 2/3 Exc | Layer 4 Exc | 160 | 0.18 nS | Distance-dependent with orientation bias |
| Layer 2/3 Exc | Layer 4 Inh | 96 | 0.35 nS | Distance-dependent with orientation bias |

**Table S1: Tabular description of the model, part 1.**

| D | Neuron and Synapse models | |
|---|---|---|
| **LGN** | | |
| Type | Leaky integrate-and-fire | |
| Subthreshold dynamics | $\tau_m \frac{dV}{dt} = -(V - E_L) + R_m I_{input}$ if $(t > t^* + \tau_{ref})$ $V(t) = V_{reset}$ else | |
| Spiking | if $V(t-) < \theta \wedge V(t+) \geq \theta$<br>1. set t* = t,<br>2. emit spike with time stamp t* | |
| Variable | Name | Value |
| V(t) | Membrane potential | — |
| $\tau_m$ | Time constant of the membrane potential | 10 ms |
| $R_m$ | Membrane resistance | 34.48 MΩ |
| $E_L$ | Leak voltage | -70 mV |
| $V_{reset}$ | Reset voltage | -70 mV |
| $I_{input}$ | Input current | — |
| $\tau_{ref}$ | Refractory period | 2 ms |
| θ | Threshold | -57 mV |
| **Layer 4 and Layer 2/3** | | |
| Type | Adaptive exponential integrate-and-fire | |
| Subthreshold dynamics | $\tau_m \frac{dV}{dt} = -(V - E_L) + \Delta_T \exp\left(\frac{V - V_T}{\Delta_T}\right) + R_m g_{exc}(E_{exc} - V) + R_m g_{inh}(E_{inh} - V) - w$ if $(t > t^* + \tau_{ref})$ $V(t) = V_{reset}$ else<br>$\tau_w \frac{dw}{dt} = a(V - E_L) - w$<br>$g_{syn}(t) = \bar{g}_{syn} e^{-(t-t^*)/\tau_{syn}}$ | |
| Spiking | if $V(t-) < \theta \wedge V(t+) \geq \theta$<br>1. set t* = t,<br>2. emit spike with time stamp t*<br>3. w = w + b | |
| Variable | Name | Value |
| V(t) | Membrane potential | — |
| w | Adaptation current | — |
| $g_{syn}(t)$ | Conductance | — |
| $g_{exc}$ | Excitatory conductance | — |
| $g_{inh}$ | Inhibitory Conductance | — |
| $\tau_m$ | Time constant of the membrane potential | 8 ms for excitatory neurons and 9 ms for inhibitory neurons |
| $R_m$ | Membrane resistance | 250 MΩ for excitatory neurons and 300 MΩ for inhibitory neurons |
| $E_L$ | Leak voltage | -80 mV for excitatory neurons and - 78 mV for inhibitory neurons |
| $V_{reset}$ | Reset voltage | -60 mV |
| $E_{exc}$ | Excitatory reversal potential | 0 mV |
| $E_{inh}$ | Inhibitory reversal potential | -80 mV |
| $\tau_{ref}$ | Refractory period | 2 ms |
| ΔT | Slope factor | 0.8 mV |
| $V_T$ | Exponential initiation threshold | -57 mV for excitatory neurons and -56 mV for inhibitory neurons |
| θ | Threshold | -40 mV |
| $\tau_w$ | Adaptation time constant | 1 ms |
| a | Subthreshold adaptation | -0.8 nS |
| b | Spike-triggered adaptation | 0.08 nA |
| $\tau_{exc}$ | Excitatory synaptic time constant | 1.5 ms |
| $\tau_{inh}$ | Inhibitory synaptic time constant | 4.2 ms |
| $\tau_{inh}$ | Inhibitory synaptic time constant | 4.2 ms |
| $g_{syn}$ | Base conductance of the synapse | — |

**Table S2: Tabular description of the model, part 2.**

| E | Plasticity | |
|---|---|---|
| Type | Short-term plasticity | |
| Model | $\frac{dx}{dt} = \frac{z}{\tau_{rec}} - ux\delta(t - t^*)$<br>$\frac{dy}{dt} = -\frac{y}{\tau_{psc}} + ux\delta(t - t^*)$<br>$\frac{dz}{dt} = \frac{y}{\tau_{psc}} - \frac{z}{\tau_{rec}}$ | |
| Variable | Name | Value |
| x | Fraction of synaptic vesicles in the readily releasable pool | — |
| y | Fraction of synaptic vesicles in the synaptic cleft | — |
| z | Fraction of inactive synaptic vesicles | — |
| t* | Spiking time | — |

| Source | Target | Effective use of the synaptic resources | Recovery time constant ($\tau_{rec}$) | Synaptic current time constant ($\tau_{psc}$) |
|---|---|---|---|---|
| LGN ON | Layer 4 Exc | 0.75 | 125 ms | 1.5 ms |
| LGN OFF | Layer 4 Exc | 0.75 | 125 ms | 1.5 ms |
| LGN ON | Layer 4 Inh | 0.75 | 125 ms | 1.5 ms |
| LGN OFF | Layer 4 Inh | 0.75 | 125 ms | 1.5 ms |
| Layer 4 Exc | Layer 4 Exc | 0.75 | 30 ms | 1.5 ms |
| Layer 4 Exc | Layer 4 Inh | 0.75 | 30 ms | 1.5 ms |
| Layer 4 Inh | Layer 4 Exc | 0.75 | 30 ms | 4.2 ms |
| Layer 4 Inh | Layer 4 Inh | 0.75 | 30 ms | 4.2 ms |
| Layer 4 Exc | Layer 2/3 Exc | 0.75 | 30 ms | 1.5 ms |
| Layer 4 Exc | Layer 2/3 Inh | 0.75 | 30 ms | 1.5 ms |
| Layer 2/3 Exc | Layer 2/3 Exc | 0.75 | 30 ms | 1.5 ms |
| Layer 2/3 Exc | Layer 2/3 Inh | 0.75 | 30 ms | 1.5 ms |
| Layer 2/3 Inh | Layer 2/3 Exc | 0.75 | 30 ms | 4.2 ms |
| Layer 2/3 Inh | Layer 2/3 Inh | 0.75 | 30 ms | 4.2 ms |
| Layer 2/3 Exc | Layer 4 Exc | 0.75 | 30 ms | 1.5 ms |
| Layer 2/3 Exc | Layer 4 Inh | 0.75 | 30 ms | 1.5 ms |

**Table S3: Tabular description of the model, part 3.**

| F | Input |
|---|---|
| Noise current | Drives LGN ON and OFF cells respectively at 17 spikes/s a 8 spikes/s when no stimulus is present |
| Visual input | $S_c(x,y) = A_c e^{\frac{-x^2-y^2}{2\sigma_c^2}}$<br>$S_s(x,y) = A_s e^{\frac{-x^2-y^2}{2\sigma_s^2}}$<br>$T(t) = K_1 \frac{(c_1(t-t_{01}))^n e^{-c_1(t-t_{01})}}{n_1^{n_1} e^{-n_1}} - K_2 \frac{(c_2(t-t_{02}))^n e^{-c_2(t-t_{02})}}{n_2^{n_2} e^{-n_2}}$<br>$rf(x,y,t) = S_c T(t) - S_s T(t-t_d)$<br>$rf_l(t) = \sum_{x=1}^{n_x} \sum_{y=1}^{n_y} rf(x,y,t)$<br>$r_l(t) = rf_l(t) \sum_{x=1}^{n_x} \sum_{y=1}^{n_y} s(x,y,t)$<br>$rf_c(x,y,t) = rf(x,y,t) - rf_l(t)$<br>$r_c(t) = \sum_{x=1}^{n_x} \sum_{y=1}^{n_y} \frac{rf_c(x,y,t) s(x,y,t)}{l_b}$<br>$I_{input}(t) = \alpha_l \frac{r_l(t)}{\beta_l + \lvert r_l(t) \rvert} + \alpha_c \frac{r_c(t)}{\beta_c + \lvert r_c(t) \rvert}$ |

| Variable | Name | Value |
|---|---|---|
| $S_c(x,y)$ | LGN cell spatial receptive field center | — |
| $S_s(x,y)$ | LGN cell spatial receptive field surround | — |
| T(t) | LGN cell temporal receptive field | — |
| rf(x,y,t) | LGN cell spatio-temporal receptive field | — |
| $r_l(t)$ | Luminance receptive field response | — |
| $r_c(t)$ | Contrast receptive field responce | — |
| $I_{input}(t)$ | Input current | — |
| $A_c$ | Spatial center Gaussian scaler | 1.0 |
| $\sigma_c$ | Spatial center Gaussian standard deviation | 0.103253 |
| $K_1$ | Positive temporal Gamma function scaler | 0.979516 |
| $t_1$ | Positive temporal Gamma function offset | -83.501129 ms |
| $c_1$ | Positive temporal Gamma function shape parameter | 1.095625 |
| $n_1$ | Positive temporal Gamma function shape parameter | 124.130728 |
| $A_s$ | Spatial surround Gaussian scaler | 0.032473 |
| $\sigma_s$ | Spatial surround Gaussian standard deviation | 0.461558 |
| $K_2$ | Negative temporal Gamma function scaler | 0.202640 |
| $t_2$ | Negative temporal Gamma function offset | -135.830829 ms |
| $c_2$ | Negative temporal Gamma function shape parameter | 0.274145 |
| $n_2$ | Negative temporal Gamma function shape parameter | 64.731311 |
| $t_d$ | Temporal offset between center and surround | 6.0 ms |
| s(x,y,t) | Stimulus time course | — |
| $n_x$ | Number of pixels in the x-axis direction | 120 |
| $n_y$ | Number of pixels in the y-axis direction | 120 |
| $l_b$ | Background luminance | 50 cd/m^2 |
| $\alpha_l$ | Luminance gain | 0.01 nA |
| $\beta_l$ | Luminance saturation parameter | 1,00E-08 cd/m^2 |
| $\alpha_c$ | Contrast gain | 0.5 nA |
| $\beta_c$ | Contrast saturation parameter | 0.0007 |

**Table S4 Tabular description of the model, part 4.**

| G | Measurements |
|---|---|
| Name | Description |
| Layer 4 Exc | Spikes: 0 neurons; Membrane potential, excitatory and inhibitory conductances: 0 neurons |
| Layer 4 Inh | Spikes: 0 neurons; Membrane potential, excitatory and inhibitory conductances: 0 neurons |
| Layer 2/3 Exc | Spikes: 38150 neurons; Membrane potential, excitatory and inhibitory conductances: 115714 neurons |
| Layer 2/3 Inh | Spikes: 18626 neurons; Membrane potential, excitatory and inhibitory conductances: 0 neurons |

**Table S5: Tabular description of the model, part 5.**

15. S. J. Van Albada, M. Helias, M. Diesmann, Scalability of Asynchronous Networks Is Limited by One-to-One Mapping between Effective Connectivity and Correlations. *PLOS*

*Comput. Biol.* **11**, e1004490 (2015).

16. C. Beaulieu, M. Colonnier, Number of neurons in individual laminae of areas 3B, 4γ, and 6aα of the cat cerebral cortex: A comparison with major visual areas. *J. Comp. Neurol.* **279**, 228–234 (1989).

17. P. Buzás, K. Kovács, A. S. Ferecskó, J. M. L. Budd, U. T. Eysel, Z. F. Kisvárday, Model-based analysis of excitatory lateral connections in the visual cortex. *J. Comp. Neurol.* **499**, 861–881 (2006).

18. A. Stepanyants, J. A. Hirsch, L. M. Martinez, Z. F. Kisvárday, A. S. Ferecskó, D. B. Chklovskii, Local Potential Connectivity in Cat Primary Visual Cortex. *Cereb. Cortex* **18**, 13–28 (2008).
